# Sustained In Vivo Perfusion of a Re-Endothelialized Tissue Engineered Kidney Graft in a Human-Scale Animal Model

**DOI:** 10.1101/2023.03.20.532642

**Authors:** Joseph S. Uzarski, Emily C. Beck, Emily E. Russell, Ethan J. Vanderslice, Matthew L. Holzner, Vikram Wadhera, Dylan Adamson, Ron Shapiro, Dominique S. Davidow, Jeff J. Ross, Sander S. Florman

## Abstract

**Introduction:** Despite progress in whole-organ decellularization and recellularization, maintaining long-term perfusion *in vivo* remains a hurdle to realizing clinical translation of bioengineered kidney grafts. The objectives for the present study were to define a threshold glucose consumption rate (GCR) that could be used to predict *in vivo* graft hemocompatibility and utilize this threshold to assess the *in vivo* performance of clinically relevant decellularized porcine kidney grafts recellularized with human umbilical vein endothelial cells (HUVECs).

**Materials and Methods:** Twenty-two porcine kidneys were decellularized and 19 were re-endothelialized using HUVECs. Functional revascularization of control decellularized (n=3) and re-endothelialized porcine kidneys (n=16) was tested using an *ex vivo* porcine blood flow model to define an appropriate metabolic glucose consumption rate (GCR) threshold above which would sustain patent blood flow. Re-endothelialized grafts (n=9) were then transplanted into immunosuppressed pigs with perfusion measured using angiography post-implant and on days 3 and 7 with 3 native kidneys used as controls. Patent recellularized kidney grafts underwent histological analysis following explant.

**Results:** The glucose consumption rate of recellularized kidney grafts reached a peak of 41.3±10.2 mg/hour at 21±5 days, at which point the grafts were determined to have sufficient histological vascular coverage with endothelial cells. Based on these results, a minimum glucose consumption rate threshold of 20 mg/hour was set. The revascularized kidneys had a mean perfusion percentage of 87.7±10.3%, 80.9±33.1%, and 68.5±38.6% post-reperfusion on Days 0, 3 and 7, respectively. The 3 native kidneys had a mean post-perfusion percentage of 98.4±1.6%. These results were not statistically significant.

**Conclusion:** This study is the first to demonstrate that human-scale bioengineered porcine kidney grafts developed via perfusion decellularization and subsequent re-endothelialization using HUVEC can maintained patency with consistent blood flow for up to 7 days *in vivo*. These results lay the foundation for future research to produce human-scale recellularized kidney grafts for transplantation.

## 1 Introduction

Every year nearly 125,000 people in the United States are diagnosed with end-stage renal disease (ESRD) and join the 800,000 Americans who suffer with the disease (United States Renal Data System, 2022). These patients require chronic dialysis or a kidney transplant for survival, although kidney transplantation is the only potential curative option. Furthermore, kidney transplantation is associated with a consistently higher 5-year survival rate compared to peritoneal or hemodialysis and a better quality of life (United States Renal Data System, 2022). However, despite its superior clinical outcomes, the shortage of donor kidney grafts limits the number of transplants that can be performed. Despite approximately 24,000 kidney transplants being performed annually, nearly 100,000 Americans remain on the waiting list (Lentine et al., 2022). Even with the expansion of living organ donation programs and extended criteria for the use of deceased donor organs, there is an insufficient supply of kidney grafts available for the increasing number of patients who are diagnosed with ESRD.

The development of a clinically viable bioengineered kidney offers the opportunity to overcome the shortage of donor organs and the need for dialysis for patients with ESRD. Unfortunately, a bioengineered kidney remains a substantial tissue engineering challenge due to this organ’s complex microarchitecture and physiology. Meaningful progress has recently been made in the directed differentiation of pluripotent stem cells into kidney organoids, and several published protocols describe production of kidney organoids containing nephrons and collecting duct structures (Miyoshi T et al., 2019; Morizane R et al., 2017; Wu H,et al., 2018; Takasato M et al., 2016) However, while organoid engineering has tremendous potential for disease modeling (Kim YK et al., 2017), diagnostic drug toxicity screening (Morizane R et al., 2015), gene editing (Kim YK et al., 2017) and other applications (Miyoshi T et al., 2019), organoids developed using these protocols lack a perfusable vasculature and a urinary drainage system. The absence of these structures limits organoid diameter to 1 to 2 cm and prevents them from recapitulating renal filtration and excretory functions, and therefore precludes their translation to clinical therapies (Miyoshi T et al., 2019).

Perfusion decellularization of intact organs enables development of a scaffold that retains the extracellular matrix architecture and composition of the vasculature and urinary drainage system that is three-dimensional (3D), acellular, and human-scale (Ott HC et al., 2008). This technology has been successfully applied to create a variety of whole-organ scaffolds including heart (Ott HC et al., 2008), lung (Ott HC et al., 2010; Petersen TH et al., 2010; Gilpin S., et al 2013), liver (Uygun BE et al., 2010; Wang Y et al, 2015), pancreas (Goh S-K et al., 2013), and kidney (Orlando G, et al., 2012; Song JJ et al., 2013; Ross EA et al., 2009; Nakayama KH et al., 2010; Bonandrini B et al., 2014; Caralt M et al., 2015; Yu Y et al., 2014; Leuning DG et al., 2019; Peloso A, et al., 2015). Promising findings have shown that decellularized kidney grafts can be recellularized and even promote differentiation of pluripotent stem cells along a renal lineage (Ross EA et al., 2009; Nakayama KH et al., 2010; Bonandrini B et al., 2014; Abolbashari M et al., 2016), however, there are limited published reports describing transplantation of these grafts in preclinical models.

The difficulty with sustaining function with decellularized animal kidneys has been reported by several authors. A study of acellular porcine kidney grafts implanted into pigs demonstrated reperfusion with sustained blood pressure without blood extravasation during surgery but complete vascular thrombosis upon histological examination at 2 weeks (Orlando G, et al., 2012). Another study found that decellularized kidney grafts were completely thrombosed 7 days after implantation into a rat model (Peloso A, et al., 2015). Separate research overcame this acute thrombogenic response using whole rat kidneys that were decellularized and then recellularized with both rat epithelial and human endothelial cells followed by perfusion in a bioreactor where they produced rudimentary filtrate *in vitro* (Song JJ et al., 2013). When implanted acutely in rats, the grafts showed no signs of thrombus formation, highlighting the importance of re-endothelializing acellular grafts. The authors did not report the length of time these grafts were studied *in vivo*. These results were also achieved in a small animal model which is on a significantly smaller scale compared to human renal grafts.

Prior to the present study, the longest reported sustained perfusion of blood in a recellularized kidney graft was an *in vitro* study performed with porcine whole blood where the grafts were reported to be patent for 24 hours (Zambron JP et al, 2018). Based on our previous research where we demonstrated that continued glucose consumption was a marker for a high rate of re-endothelization in decellularized porcine liver grafts (Mao SA et al., 2017), we hypothesized that glucose consumption by the recellularized kidney graft would also serve as a marker for its extent of endothelialization and indicate the ability of a recellularized porcine kidney graft to remain patent.

The objectives for the present study were to define a threshold glucose consumption rate (GCR) that could be used to predict *in vivo* graft hemocompatibility and then utilize this threshold to assess the *in vivo* performance of clinically relevant decellularized porcine kidney grafts recellularized with human endothelial cells in an orthotopic transplantation model.

## 2 Methods

The present study was carried out in the facilities of American Preclinical Services (Minneapolis, MN), an Association for Assessment and Accreditation of Laboratory Animal Care (AAALAC) approved facility. The study protocol was reviewed and approved by the facility’s Institutional Animal Care and Use Committee (IACUC). All animals were housed, fed, cared for, and euthanized by study personnel and animal husbandry staff of American Preclinical Services.

### 2.1 Donor kidney selection and recovery

Twenty-two porcine kidneys weighing 250 to 300 grams were obtained from adult (approximately 6 months old) male and female Landrace/Yorkshire/Duroc crossbreed pigs purchased from Midwest Research Swine (Glencoe, MN). After recovery, the kidneys were rinsed in saline, bagged, and transported on ice to a separate facility for processing.

### 2.2 Decellularization

Prior to decellularization, the kidney pairs were removed from ice and passed into a class 10,000 clean room where they were disinfected with peracetic acid (PAA) before and after cannulation. The renal artery, renal vein, and ureter on each kidney were cannulated using appropriately sized polypropylene cannulae (Value Plastics, Loveland, CO) and secured with 3-0 nylon sutures (eSutures, Mokena, IL). Cannulated kidneys were perfusion decellularized using Triton X-100 for 3 hours followed by 0.3% sodium dodecyl sulfate (SDS) overnight. The kidneys were then flushed with phosphate buffered saline (PBS) and underwent further disinfection using 1000 ppm peracetic acid prior to a final PBS flush and packaging. Perfusion pressure was held at a constant value (renal artery: 60 mmHg; renal vein: 40 mmHg; ureter: 20 mmHg) by altering the volumetric flow rate of the peristaltic pump. Perfusion alternated between the renal artery, vein, and ureter at evenly spaced intervals during the PAA and PBS stages. The decellularized kidney scaffolds were then packaged with PBS and stored at 4°C for up to 6 months.

### 2.3 Endothelial culture and seeding

To establish thromboresistance for the porcine extracellular matrix and prepare the decellularized kidney grafts for transplantation, human umbilical vein endothelial cell (HUVEC) suspensions were infused sequentially through the renal vein and artery using a perfusion bioreactor system (Figure 1). HUVECs were chosen for kidney revascularization due to the ability to create an HLA bank, favorable proliferation kinetics, phenotypic plasticity with the ability to form fenestrations in vitro (Hamilton RD et al., 2007), and past success with HUVECs for liver recellularization (Shaheen et al., 2020). The bioreactor system used custom perfusion software and a peristaltic pump to drive the flow of culture media into the kidney through either the renal vein or artery, while maintaining the volumetric flow rate or pressure at a specified value. This system enabled more consistent recellularization among different donor kidneys and across different recellularization lots by closely regulating environmental parameters (e.g., perfusion pressure/flow, temperature, pH, and oxygen tension) during seeding and subsequent perfusion culture.

**Figure 1.**
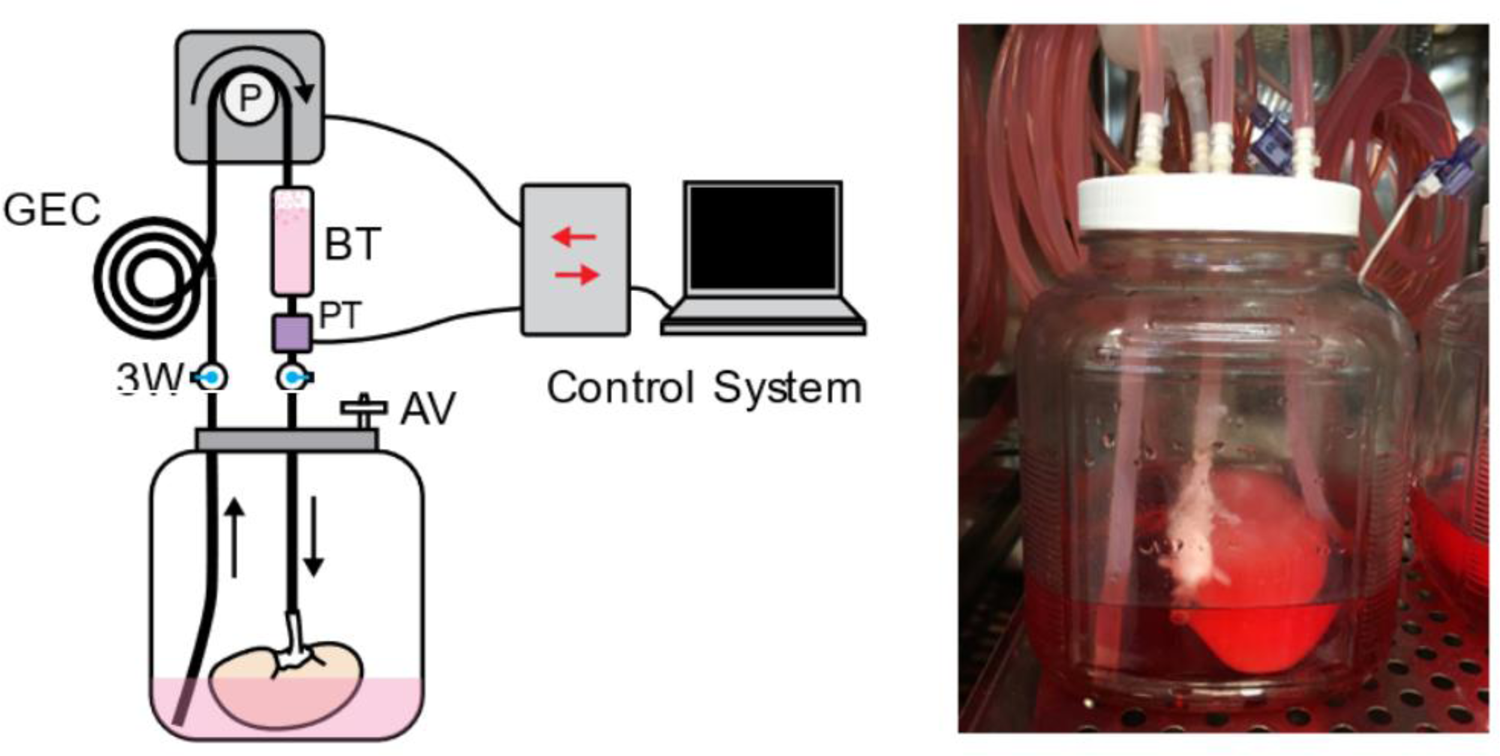
The perfusion system used for kidney recellularization. The system consisted of a reservoir holding the kidney, an air vent (AV), a pressure transducer (PT) to monitor perfusion pressure, a gas exchange coil (GEC) to allow for diffusion of gas into the perfusate, a bubble trap (BT), and several three-way (3W) stopcocks to direct the flow of the media to the kidney. Perfusion was driven by a peristaltic pump under control by custom perfusion software.

The decellularized kidneys were mounted in the sterile bioreactors inside humidified incubators. They were then perfused through the renal vein at 100 mL/min with a pressure less than 20 mmHg in custom antibiotic-free endothelial media containing endothelial basemedia (R&D Systems, Minneapolis, MN, #CUST01707), sodium bicarbonate (Sigma-Aldrich, St. Louis, MO, #S5761), fetal bovine serum (FBS) (Fisher Scientific), ascorbic acid (Amresco, Solon, OH, #0764), hydrocortisone (Tocris, Minneapolis, MN, #4093), fibroblast growth factor (FGF) (R&D Systems, #233-FB/CF), vascular endothelial growth factor (VEGF) (R&D Systems, #293-FB/CF), epidermal growth factor (EGF) (R&D Systems, #236-EG), recombinant insulin-like growth factor (R3-IGF) (Sigma Aldrich, #85580C), heparin (Sigma Aldrich, #H3393), and acetic acid (Sigma Aldrich, #A6283)) for at least 3 days to confirm sterility. The media was replaced with endothelial media containing 1% penicillin/streptomycin prior to seeding. Primary cryopreserved HUVECs were expanded in 5-chamber CellSTACK cell culture chambers (Corning, Glendale, AZ) using antibiotic-free endothelial media and were lifted for seeding at passage 5-8 (Lonza, Walkersville, MD). After lifting, HUVECs were passed through a 70 µm strainer and were resuspended at a concentration of 1 million cells/mL. Cells were seeded through the vein with 100 mL cell suspension without perfusion for 1 hour and then 50 mL cell suspension was injected into the venous perfusion flow field at 50 mL/min. After 24 hours, perfusion was switched from the vein to the artery, the media was changed, and another 150 million HUVEC cells were seeded through the artery using the same protocol. At 24 hours after the first arterial seeding, the media was replaced, and the flow was increased to 100 mL/min. Metabolite levels in bioreactor media samples were measured daily using a BioProfile FLEX Analyzer (Nova Biomedical, Waltham, MA). Media was changed on alternating days and GCR was monitored. Between 1 to 6 days after the first arterial seeding, the kidney grafts were seeded through the artery a second time using the previously described arterial seeding protocol. The media was changed 24 hours after the serial arterial seeding and then the frequency and volume of media changes was adjusted as needed to prevent total glucose depletion.

### 2.4 Cell metabolism

Cell metabolic activity was monitored daily by analyzing a bioreactor media sample. Sample metabolite concentrations were obtained from a Cedex Bio HT Analyzer (Roche) to monitor metabolism of glucose, glutamine, glutamate, ammonia, and lactate dehydrogenase (LDH). When glucose levels fell below 0.2 g/L or ammonia levels rose above 1 mM, additional media was added to the bioreactor reservoir.

### 2.5 Acute ex vivo transplant model

Functional revascularization of control decellularized porcine kidneys (n=3) and re-endothelialized kidneys (n=4) was tested using an acute *ex vivo* transplant model. This enabled perfusion of the kidneys under physiological hemodynamic conditions in an observable setting (Figure 2). This blood loop system was used for evaluation of vascular integrity, consistency of tissue perfusion, and distribution of flow across the renal vasculature. Catheters were placed into the carotid artery and jugular vein in anesthetized adult Yorkshire Cross pigs receiving intravenous heparin injections to maintain activated clotting time between 175 to 225 seconds. Tygon tubing and barbed Luer adapters were used to create a loop linking the carotid artery and jugular vein. A Transonic flow meter and 8 mm flow probe (T402 modular research console, Transonic Systems Inc., Ithaca, NY) and pressure T were integrated in the system downstream from carotid access and upstream from the graft. Baseline flow and graft inflow were recorded, and angiographies performed over time.

**Figure 2.**
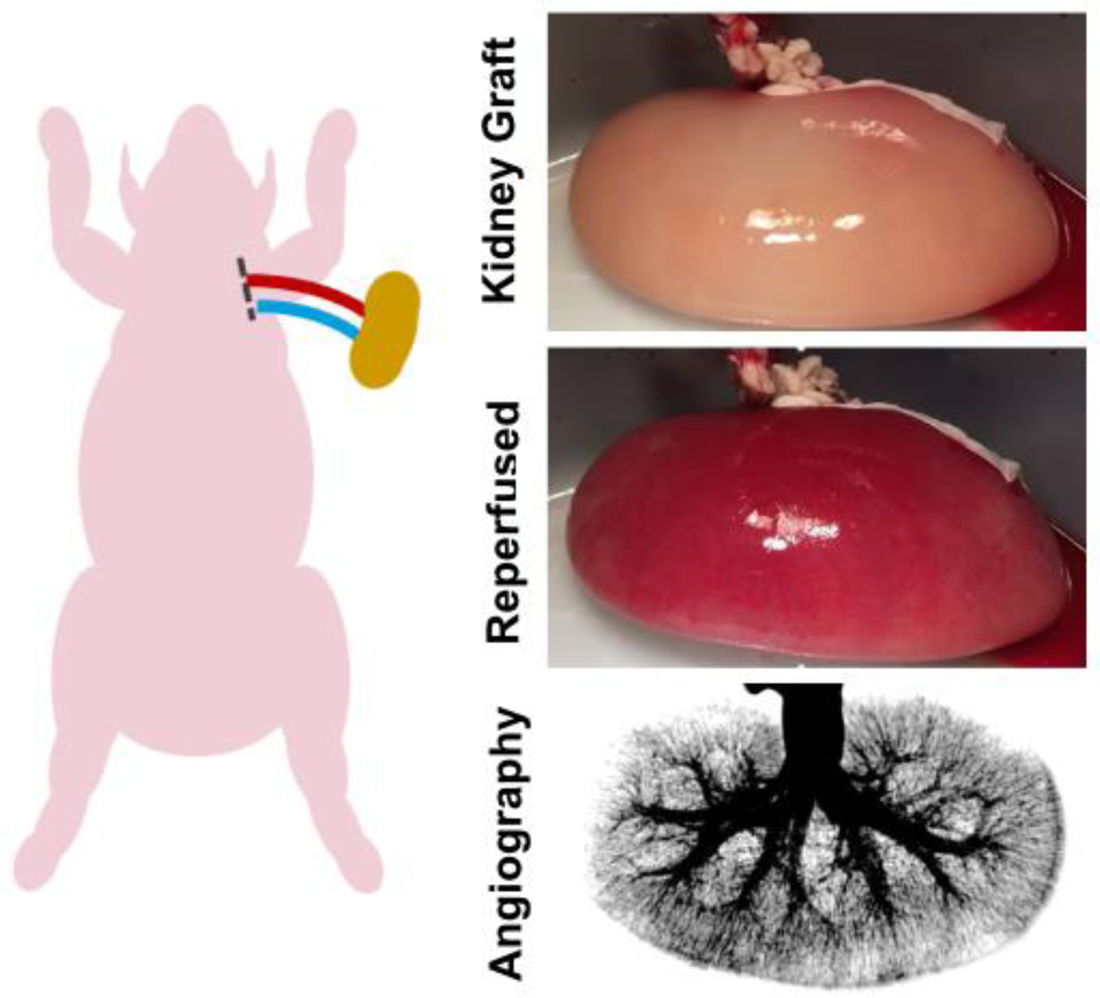
Ex vivo transplant model. An *ex vivo* blood loop was created by placing catheters in the carotid artery and jugular vein of an anesthetized adult pig. The catheter lines were connected via Luer adapters to the renal artery and vein of the endothelialized kidney graft.

### 2.6 Bioengineered kidney graft implantation

Following confirmation of functional patency using the *ex vivo* transplant model, functional revascularization was evaluated in a porcine orthotopic kidney transplantation model. Although heterotopic kidney transplantation is the clinical standard, an orthotopic model was chosen here to create a larger surgical field for placement of the endothelialized kidney grafts and to enable graft repositioning to ensure sufficient renal blood flow.

Nine kidney grafts were implanted in separate Yorkshire Cross pigs weighing 75 to 95 kg. Implantation occurred 19 to 29 days after initial HUVEC seeding depending on achievement of the targeted GCR (Table 1). Prior to implantation, the kidney grafts were flushed with heparinized saline and the ureter was ligated. Each kidney was spray coated with 2 mL Tisseel (Baxter, Deerfield, IL) using the EASYSPRAY Set (Baxter) and the Tisseel was allowed to set for 3 minutes. Under general anesthesia and heparin administered to raise the activated clotting time to 175 to 225 seconds, the kidney grafts were implanted into pigs. Following a laparotomy, surgeons performed a splenectomy to mitigate acute rejection of the HUVECs, and a nephrectomy. The revascularized grafts were anastomosed to the renal vein and artery or by a side anastomosis to the abdominal aorta and inferior vena cava. The graft was secured to the abdominal wall using a biological sling. MIROMESH Biological Matrix (Miromatrix Medical, Eden Prairie, MN) was used as the sling material in the first 3 cases before switching to the native peritoneum to obtain better stability in remaining cases after graft movement and thrombosis was observed. Vital signs and arterial flow to the kidney were monitored using the Transonic flow meter until the flow stabilized.

**Table 1.** Surgical Details and Outcomes

| **Animal #** | **Arterial Anastomosis** | **Venous Anastomosis** | **Sling Material** | **Clopidogrel & Aspirin Administered** | **Days Between Arterial HUVEC Seedings** | **Peak GCR (mg/hr)** | **End GCR (mg/hr)** | **Days of 3D Culture Before Implant** | **Patent at POD 7?** |
| --- | --- | --- | --- | --- | --- | --- | --- | --- | --- |
| 1 | Renal artery | Renal vein | MIROMESH | No | 6 | 39.54 | 34.16 | 22 | Yes |
| 2 | Renal artery | Renal vein | MIROMESH | No | 2 | 29.95 | 29.95 | 27 | No - Thrombosed |
| 3 | Renal artery | Renal vein | MIROMESH | No | 5 | 36.51 | 36.51 | 29 | No - Thrombosed |
| 4 | Abdominal aorta | Renal vein | MIROMESH | No | 1 | 41.01 | 41.01 | 21 | No - Terminated POD0 due to uncontrolled graft bleeding |
| 5 | Renal artery | Renal vein | Peritoneum | Daily starting POD1 | 1 | 30.25 | 27.09 | 29 | Yes |
| 6 | Renal artery | Renal vein | Peritoneum | Daily starting POD1 | 2 | 37.92 | 31.91 | 28 | Yes |
| 7 | Renal artery | Inferior vena cava | Peritoneum | Daily starting POD1 | 1 | 33.67 | 33.66 | 28 | Yes |
| 8 | Renal artery | Renal vein | Peritoneum | Daily starting POD1 | 2 | 43.00 | 41.20 | 25 | No - Thrombosed |
| 9 | Abdominal aorta | Renal vein | None (graft implanted retroperitoneally) | Daily starting POD1 | 1 | 41.91 | 36.20 | 19 | Yes |

GCR = glucose consumption rate; POD = post-operative day

The recellularized implanted kidney grafts were monitored with angiography post-operatively and at days 3 and 7. Two of the native kidneys were also monitored at similar timepoints. Daily medications administered included aspirin and ketoprofen for pain management, ceftiofur sodium and ceftiofur crystalline free acid for infection prophylaxis, and methylprednisolone to mitigate immunological rejection of HUVECs. After observing localized vascular thrombosis at the anastomosis site in 2 implanted grafts at days 3 and 7, daily clopidogrel anticoagulation therapy was administered to subsequent recipient pigs starting on post-operative day 1.

### 2.7 Angiography

Angiography was used to assess kidney graft patency following implantation. A sheath was placed in the carotid or femoral artery and a catheter advanced to the location of arterial blood supply of the graft. Omnipaque 300 contrast agent (GE Healthcare, Chicago, IL) was then injected via the catheter. The contrast agent was injected with subtracted angiography using the Pie Medical Software on the Siemens Leonardo Workstation with a Siemens Artis instrument, and images were recorded until the contrast had completely cleared from the kidney. Angiography images were acquired at the frame at which the maximum loading of contrast was retained in the kidney. Using ImageJ, a grayscale histogram of pixels was obtained for the region containing the kidney and the pixels with the same grayscale value as the background surrounding the kidney were subtracted. Percent perfusion was then calculated as the percentage of remaining number of pixels retained within the kidney histogram compared to the original number of pixels prior to background subtraction. Results are presented as percent graft perfusion. Following the same procedure, contralateral native kidneys were analyzed in 3 of the recipient pigs to compare to the patency of the kidney grafts.

### 2.8 Histology

Histologic samples were embedded in paraffin, sectioned at 5 µm thickness, and were stained with hematoxylin and eosin (H&E) and Masson’s Trichrome staining procedures (Scientific Solutions, LLC). Imaging was performed using Zeiss Zen software and Axiocam 105 color brightfield microscope camera. For explanted patent kidney grafts, the pigs were euthanized with intravenous barbiturate and the graft was excised and flushed with 300 mL PBS followed by 300 mL 10% neutral buffered formalin (NBF, VWR). If the kidneys could not be fully flushed due to thrombosis, they were cut into 1 cm slices and submerged in 10% NBF to enable further processing similar to the patent kidneys. Regions (1 cm thick) were cut from the upper and lower pole as well as the midline of the kidney and were submerged in 10% NBF for 2 days.

### 2.9 Immunofluorescence Staining

Paraffin embedded sections (Scientific Solutions, Fridlay, MN) were deparaffinized, rehydrated, and subjected to antigen retrieval in a Decloaking Chamber NxGen (Biocare Medical, Pacheco, CA). Slides were washed with sodium borohydride in PBS to reduce autofluorescent signal from red blood cells. They were then co-stained with appropriate primary and secondary antibodies and DAPI, then mounted using ProLong Diamond Antifade Mountant (Thermo Fisher Scientific). Imaging was performed using Jenoptik Gryphax software and Progres Gryphax fluorescent microscope camera. Anti-CD31 antibody (Abcam, Freemont, CA, #ab28364) is reactive to both human and pig endothelial cells. Anti-CD31 antibody (Abcam, #ab187377) is reactive to only human endothelial cells. A co-stain with these antibodies was used to differentiate between HUVEC and recipient pig endothelial cells. Anti-VE-cadherin antibody (Abcam, # ab33168) was also used to identify both human and pig endothelial cells. Anti-human nucleoli antibody (Abcam, # ab190710) was used to identify human endothelial cells. Anti-Collagen I antibody (Abcam, #ab34710), Anti-Collagen IV antibody (Abcam, #ab6586), and Anti-Laminin antibody (Abcam, #ab11575) were used for matrix characterization. AlexaFluor 488 (Thermo, #A11034, #A11029) and 555 secondary antibodies (Thermo, #A21429, #A21424) were used to detect the primary antibodies.

### 2.10 Statistics

A Kruskal-Wallis ANOVA was performed on the angiography data using GraphPad Prism 8.1.1 software (GraphPad Software, Inc., La Jolla, CA), where p ≤ 0.05 was considered significant. Data are reported as mean ± standard deviation.

## 3 Results

The kidneys obtained from adult donor pigs retained their native gross shape, size, and ECM proteins, including collagen I, laminin, and collagen IV following decellularization (Figures 3A and 3B). The resulting matrix retained the appropriate 3D architecture for renal microstructures, including blood vessels, glomeruli, nephron tubules, collecting ducts, and papillae (Figure 3C). Sixteen kidneys were used for the study to determine the GCR threshold and 9 were used to assess the vascular patency of implanted re-endothelialized kidney grafts.

**Figure 3.**
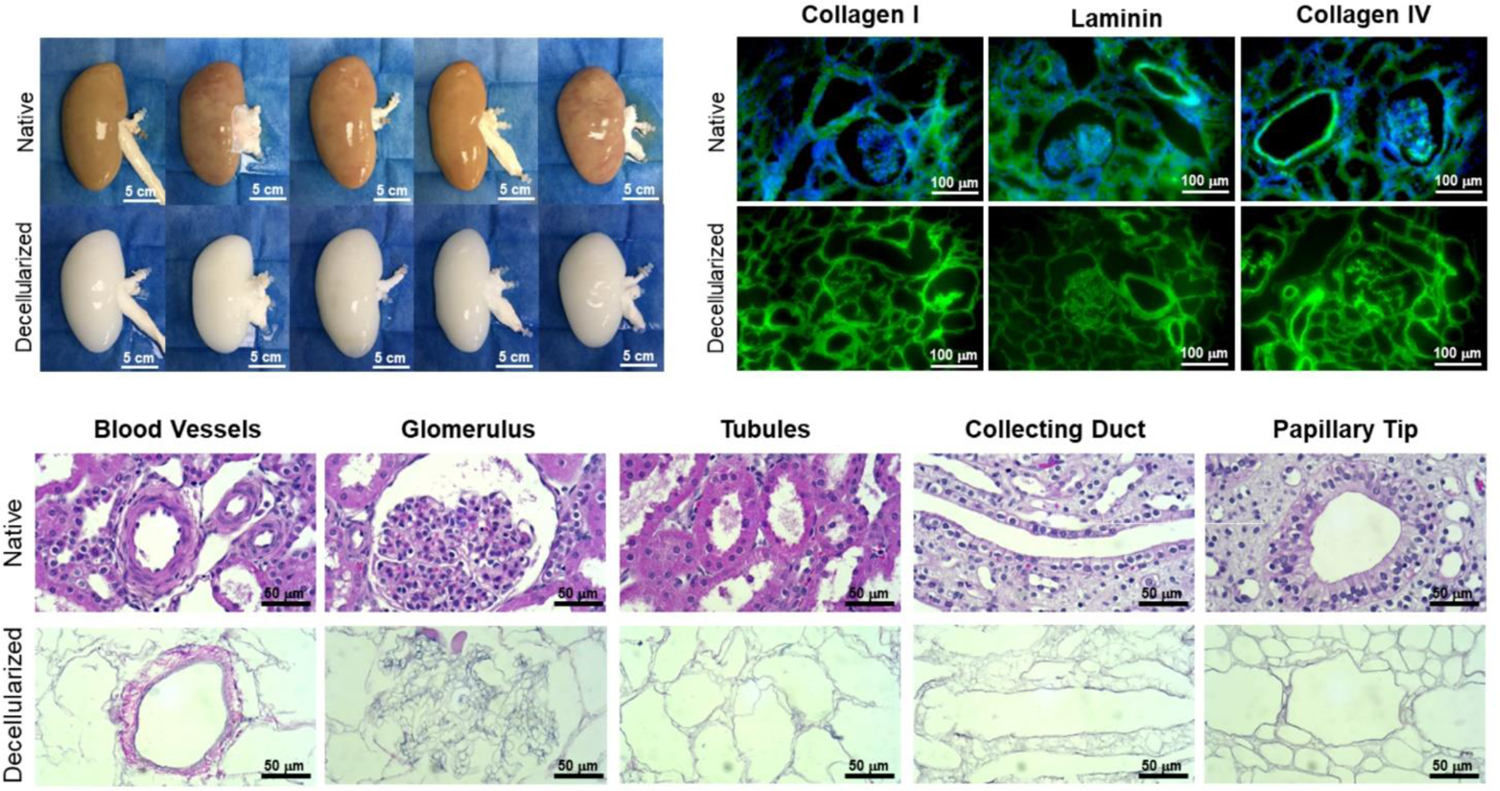
Representative gross, immunofluorescence, and histological images comparing native and decellularized kidneys. **(A)** Decellularization caused a gross loss of color as the native porcine cells were solubilized and extracted, while the original size, shape, and structure of the kidney were well-retained in the decellularized scaffold (scale bars, 5 cm). **(B)** Immunofluorescence staining showed retention of collagen I, laminin, and collagen IV (green) and complete removal of nuclei (blue DAPI stain) (scale bars, 100 µm). **(C)** High-magnification H&E images revealed the ECM structures left behind in decellularized kidneys, including blood vessels, glomeruli, nephron tubules, collecting ducts, and papillae (scale bars, 50 µm)

### 3.1 Determination of threshold glucose consumption rate (GCR)

Daily monitoring of total glucose consumption rate GCR by endothelialized grafts demonstrated GCR was a non-destructive indicator of re-endothelialization. While GCR remained low on days 0 to 7 after seeding, after 7 to 14 days of perfusion culture the GCR gradually increased with a peak mean GCR achieved at 21±5 days (41.3±10.2 mg/hour) (Figure 4A). After continued culture, the GCR plateaued and eventually started to decline following confluency.

**Figure 4.**
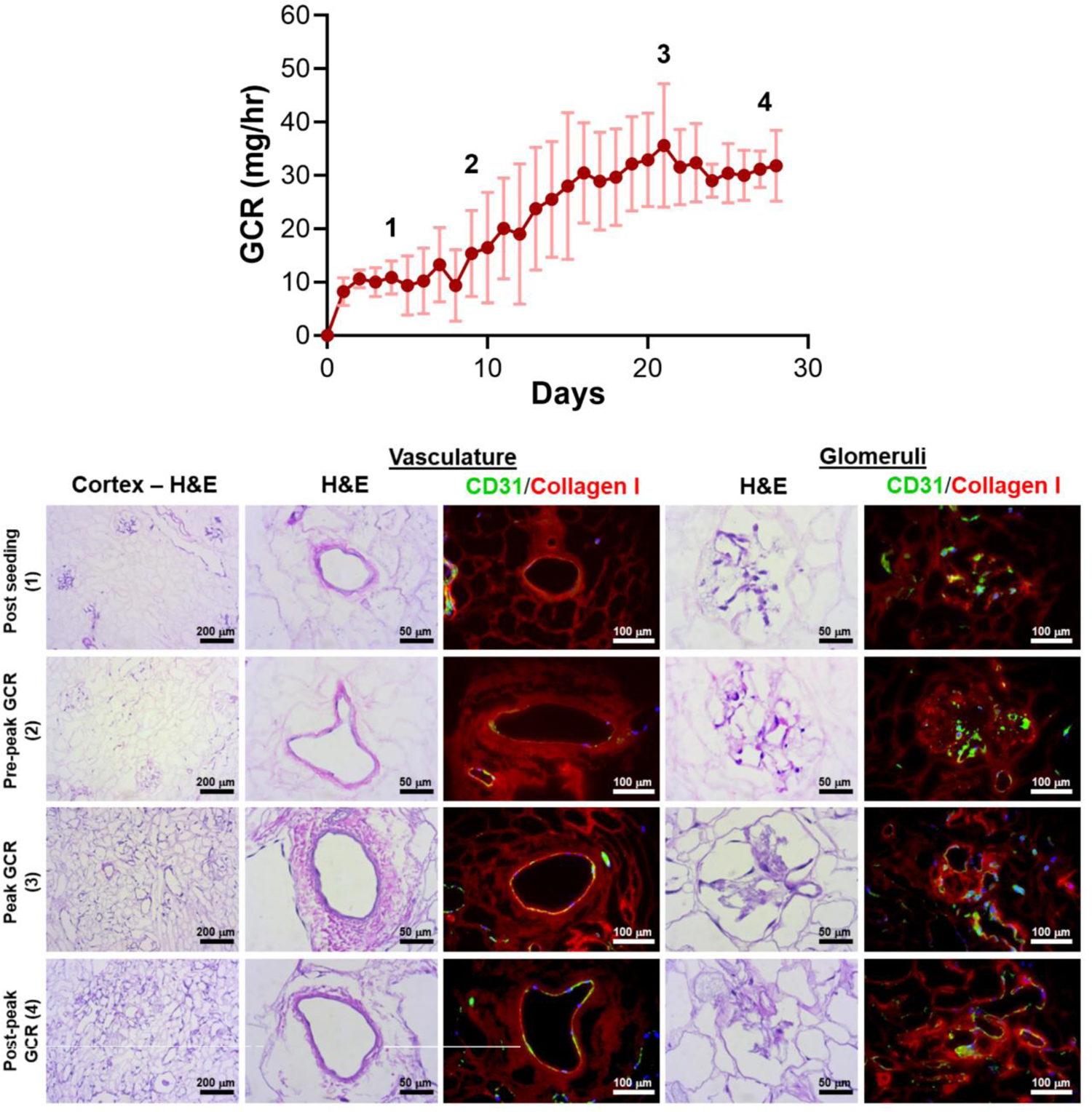
Correlation of glucose consumption kinetics with vascular endothelial cell coverage for decellularized porcine kidneys reendothelialized with HUVECs. **(A)** Total glucose consumption rate (GCR) by endothelial cells was calculated by monitoring the changes in glucose concentration in bioreactor media samples over time. Four distinct zones (1 through 4) were defined to distinguish endothelial cell coverage based on GCR and culture time. **(B)** Representative H&E images and immunofluorescence images revealed endothelialization of the vasculature and glomerular capillaries at each zone (green: CD31 staining HUVECs, red: collagen I staining the porcine matrix; blue: DAPI staining cell nuclei).

HUVECs were found to engraft both large vessels and glomerular capillaries early after seeding (Figure 4B, “Post Seeding (1)”). The density of endothelial cells in blood vessels and glomeruli increased between days 7 to 14 (Figure 4B “Pre-peak GCR (2)”). The native porcine vasculature was uniformly reconstituted with HUVECs with all histologic sections evaluated showing complete circumferential coverage when peak mean GCR was achieved (Figure 4B, “Peak GCR (3)”). In this peak period, the endothelial cells were evident in large blood vessels and glomerular capillaries.

Measurement of perfusion using the acute *ex vivo* transplant model showed the 3 control decellularized kidneys thrombosed rapidly within 5 minutes of being perfused with the inlet volumetric flow rate rapidly declining to zero (Figures 5A and 5B). Similar results were seen for the single endothelialized kidney that had a GCR below 20 mg/hour (Figures 5A and 5B) which was associated with incomplete vascular coverage (Figure 4B, “Post Seeding (1)”). In contrast to the above, the 3 endothelialized kidneys expressing Peak GCRs greater than 20 mg/hour had sustained, consistent perfusion rates of over 100 mL/min after 30 minutes of continuous blood perfusion (89.9 ± 5.7% of baseline values). Quantification via angiography stills after a minimum of 30 minutes confirmed consistently high perfusion percentage of endothelialized grafts (Figure 5B).

**Figure 5.**
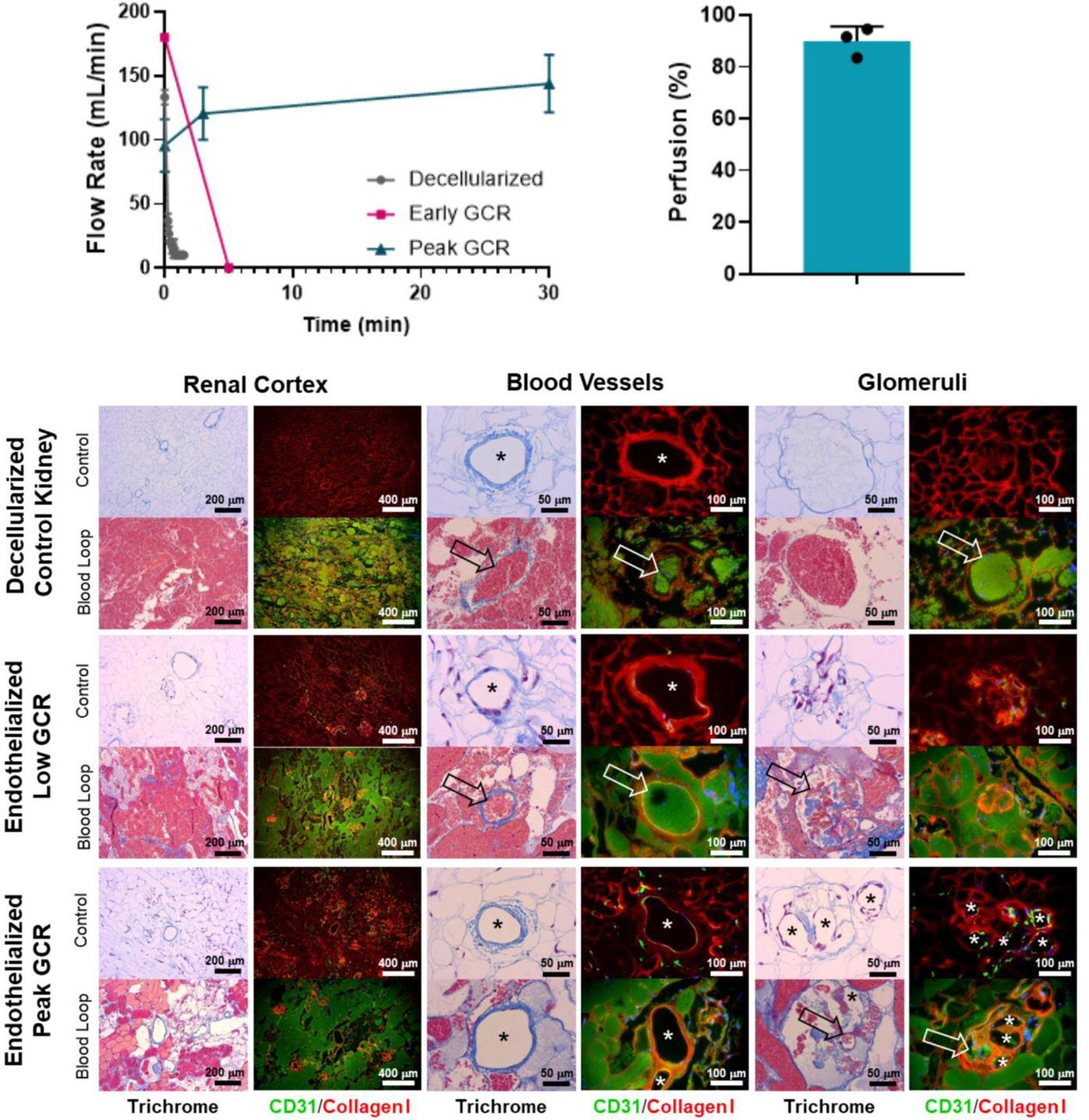
A Transonic flow meter was used to monitor volumetric flow rate of blood leading into the renal artery of the organ. Decellularized kidneys (no cells, n=3) and an endothelialized graft with Low GCR (n=1) did not maintain blood flow beyond several minutes. Endothelialized grafts near Peak GCR maintained consistent perfusion throughout the entire study period (30 to 80 minutes, n=3). Quantified angiographic images showed a high percentage of graft perfusion. Data for perfusion was reported as mean + standard deviation and with individually plotted data points. (**C**) Representative Masson’s Trichrome stain or CD31/Collagen I immunofluorescence images (asterisks denote the lumens of patent blood vessels; open arrows point to thrombosed blood vessels).

Histologically, the vasculature in grafts at Peak GCR remained patent and free of thrombus throughout the recellularized kidney after perfusion with blood (Figure 5C). Based on these results, it was determined that a minimum glucose consumption rate threshold of 20 mg/hour was required for endothelialized grafts to maintain consistent perfusion in acute (>30 minutes) *ex vivo* blood loops.

### 3.2 Vascular Patency of re-endothelialized kidney grafts

Table 1 reviews the outcomes of the 9 implanted kidney grafts. Reperfusion was initially uniform for all revascularized kidneys with contrast reaching all the way to the cortical edges of the kidney (Figure 6B, Supplementary Movie 1). One pig died unexpectedly due to excessive peri-operative bleeding and 2 pigs were terminated at days 3 and 7 due to lack of perfusion resulting from thrombosis (Figure 6C) which was suspected to be a result of graft movement and vessel torsion.

**Figure 6.**
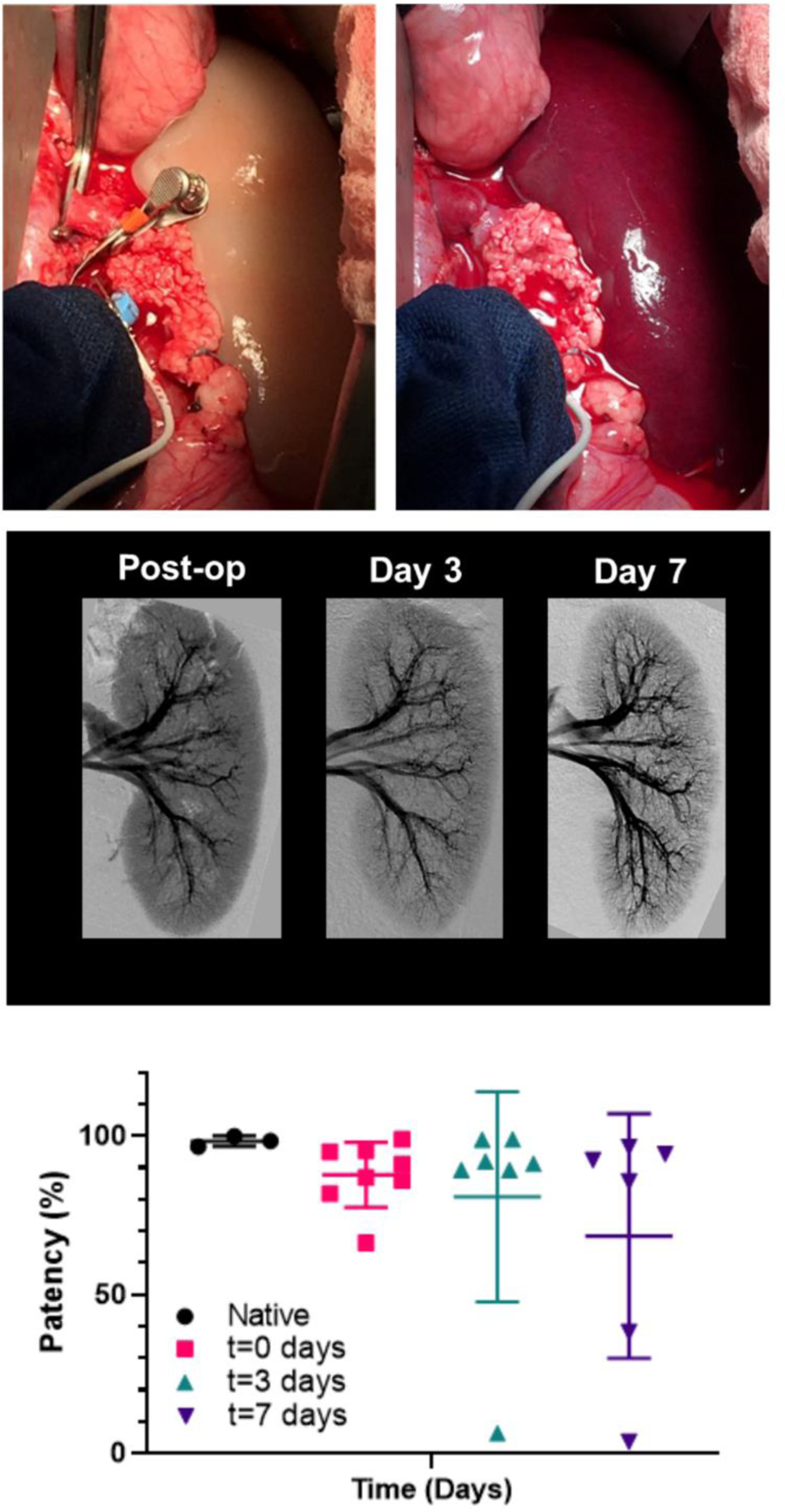
Perfusion of endothelialized kidney grafts. **(A)** Anastomosed kidney graft before and after reperfusion. (**B**) Representative angiography of kidney graft through post-operative day 7. (**C**) Percent perfusion quantified from angiographies performed on the day of surgery through post-operative day 7 for implanted recellularized kidneys (n=9) and control contralateral native kidneys (n=3). Lines represent medians and interquartile ranges.

Five of the remaining 6 grafts (83.3%) remained patent through 7 days after implantation showing renal perfusion during follow-up angiography. All pigs which developed thrombosis had kidney grafts stabilized by MIROMESH and were not administered clopidogrel or aspirin. Vascular patency was observed in one implant at day 7 in the absence of anti-coagulant therapy before clopidogrel was added to address thrombosis at the anastomoses. The 3 native kidneys had a mean perfusion percentage of 98.4 ± 1.6% compared to 87.7 ± 10.3%, 80.9 ± 33.1%, and 68.5 ± 38.6% for the revascularized kidneys post-reperfusion, on day 3, and on day 7 respectively. These differences were not statistically significant.

### 3.3 Histologic Findings for re-endothelialized kidney grafts

Patent grafts were explanted for histological analysis at approximately 2 hours post-reperfusion (acute) and at Days 3 and 7. The grafts were flushed with saline through the renal artery to remove residual blood before fixation. Histological trichrome staining of the acute explanted kidney showed the vasculature, including glomerular capillaries, remained clear of thrombus (Figure 7). The nephron tubules and surrounding interstitium showed accumulation of clotted blood due to residual vascular leakage that could not drain through the ligated ureter. Glomerular capillaries in grafts explanted at subsequent follow-up time points (Days 3 and 7) showed evidence of occlusion associated with a lack of endothelial lining (Figure 7).

**Figure 7.**
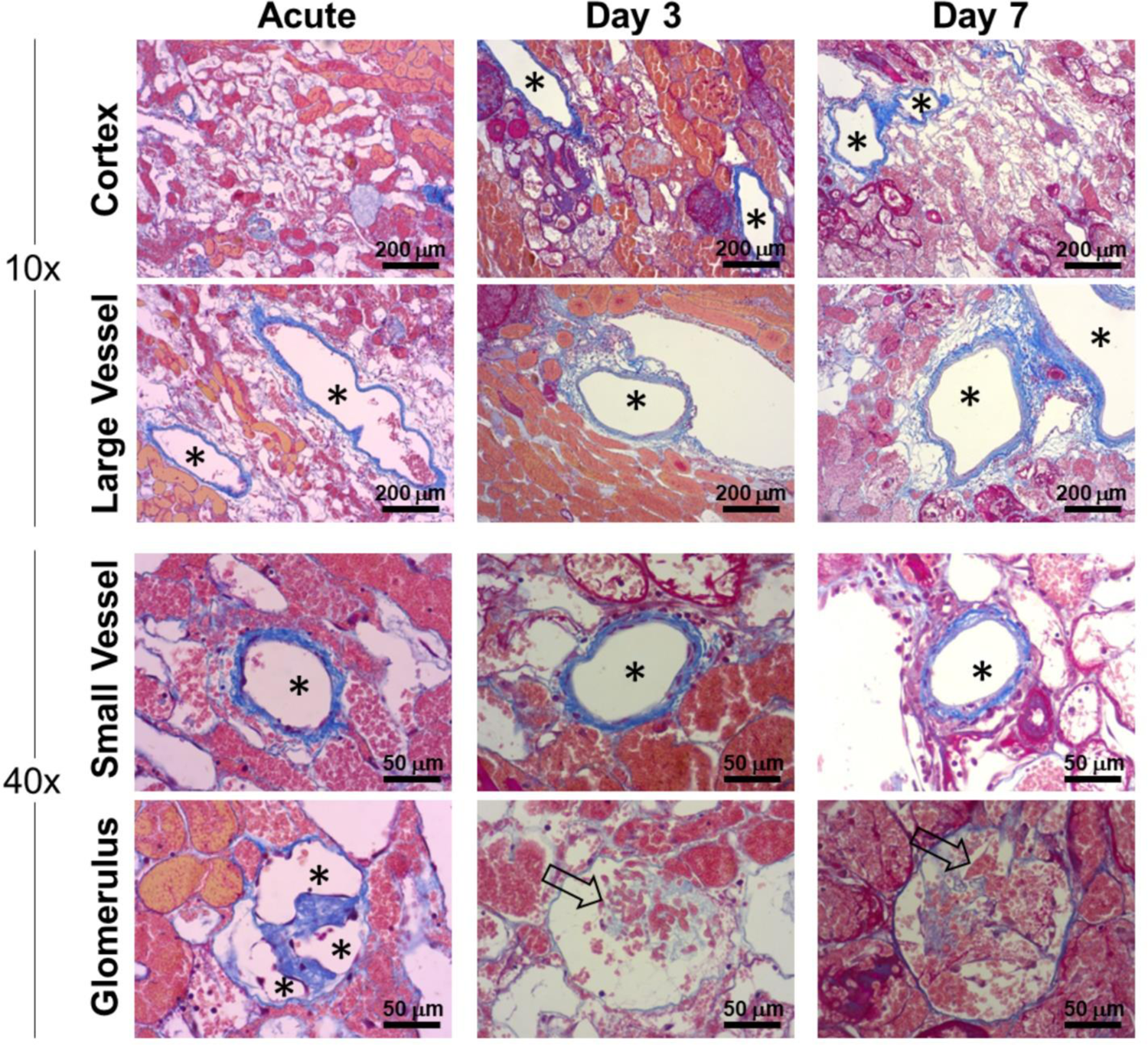
Histological characterization of vascular patency in implanted endothelialized kidney grafts acutely (approximately 2 hours post-reperfusion), Day 3 and Day 7. Representative trichrome images show large (>250 µm) and small (<100 µm) blood vessels that remained patent and clear of residual blood after post-explant flushing. Asterisks denote the lumens of patent blood vessels. Open arrows indicate occluded glomerular capillaries.

Despite methylprednisolone administration to limit immune rejection of human cells, the HUVECs were observed to gradually disappear from the kidney grafts between post-operative day 3 and 7, and grafts explanted from day 7 were absent of any HUVECs as determined by immunofluorescence staining of fixed tissue (Figure 8A and 8B; Supplementary Figures 1 and 2). Despite this apparent lack of viable human endothelial cells, the vasculature remained patent and the luminal surface did not initiate a clotting cascade, as evidenced by the lack of thrombus (Figures 7 and 8). By Day 7 porcine vascular endothelial cells were observed within the renal vasculature starting with the minor blood vessels (Figure 8B).

**Figure 8.**
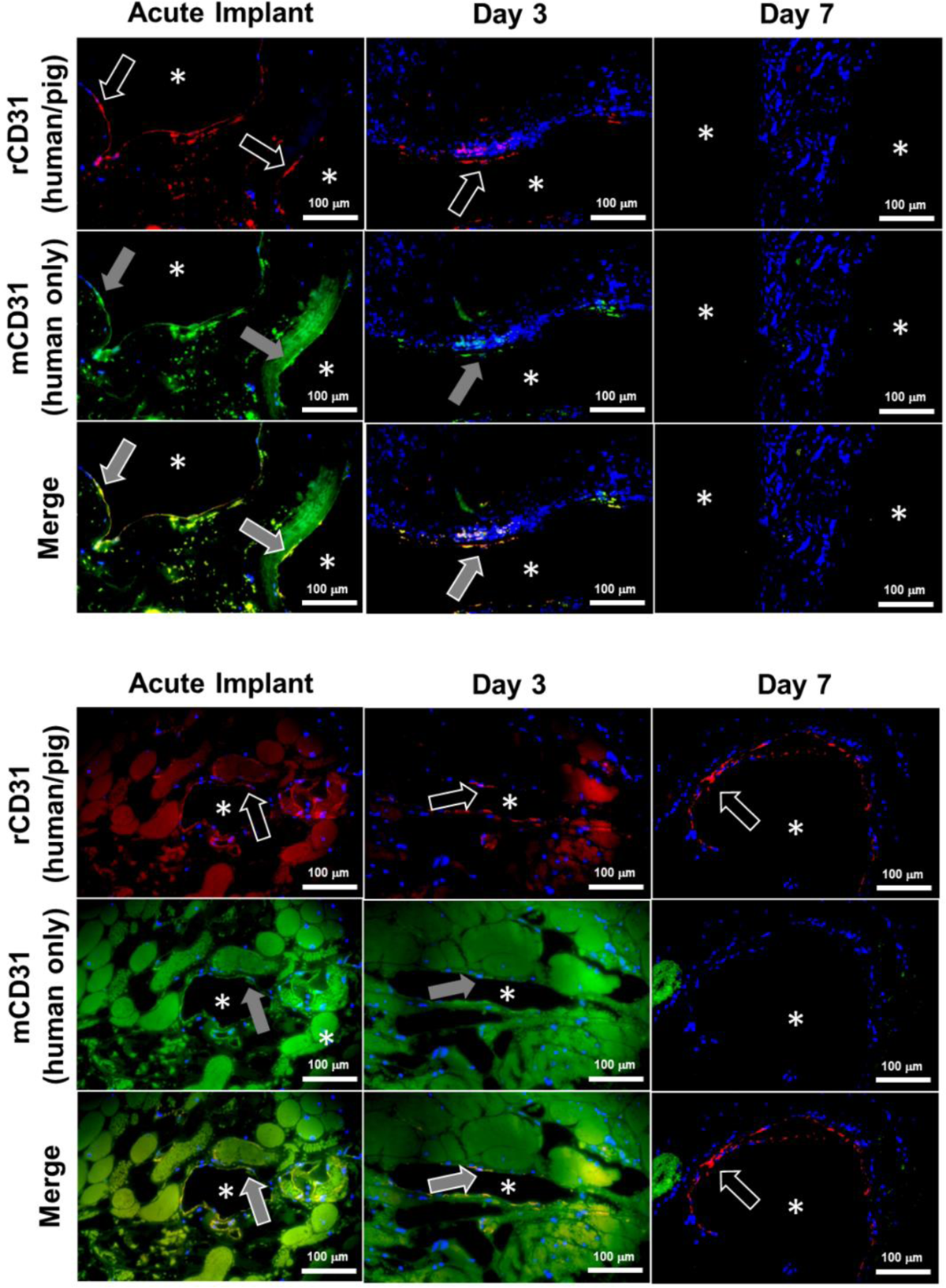
Donor human endothelial cell turnover with recipient porcine endothelial cells in chronically transplanted endothelialized kidney grafts. Endothelial cells were identified as donor (human) or recipient (porcine) in explanted kidney grafts using immunofluorescence staining. Images show the luminal surfaces of **(A)** major blood vessels exceeding 500 µm in diameter, or **(B)** minor blood vessels less than 250 µm in diameter. Asterisks denote the lumens of patent blood vessels. White arrow outline indicates positive reactivity by rabbit CD31 antibody (rCD31; red stain), which reacts to both human and pig endothelial cells. Grey arrow fill indicates positive expression by mouse CD31 antibody (mCD31; green stain), which reacts positively only in human endothelial cells. Top row: rCD31/DAPI overlay. Middle row: mCD31/DAPI overlay. Bottom row: rCD31/mCD31/DAPI merged images. Orange shows colocalization of both antibodies indicating human endothelial cells. Note that the presence of residual blood causes faint background staining in some images. Donor human endothelial cells were completely absent from the graft by Day 7,while new pig endothelialization started to occur at Day 7. Scale bars, 100 µm.

## 4 Discussion

Perfusion decellularization technology enables the removal of cells and cellular antigens from kidneys with the promise to serve as the scaffold for bioengineering of functional human-scale kidneys and thus, eliminate the need for dialysis and the transplant waiting list (Song JJ et al.,2013; Uzarski JS et al., 2014; Ott HC et al.,2008). However, an essential design requirement for a transplantable bioengineered organ is a functional vascular supply that can maintain physiological blood flow without thrombosing. Decellularized organs by definition do not meet this requirement due to widespread exposure of the vascular basement membrane to blood, the proteins of which are highly thrombogenic (Balleisen L et al., 1980; Willette RN et al., 1994) and initiate rapid thrombosis even in an animal treated with heparin (Bonandrini B et al., 2014; Mao SA et al.,2017) (Figure 5). Therefore, to create a foundation for eventual transplantation of parenchymal-seeded grafts, this study was conducted to first focus on endothelialization of the renal vasculature of decellularized porcine kidneys with human vascular cells and demonstrate sustained perfusion.

The renal vasculature branches from the renal artery and vein into interlobar vessels down to individual glomerular capillaries and the vasa recta, which are essential to renal function by facilitating reabsorption/excretion via proximity to the intertwined network of nephron tubules (Molema G et al., 2012). Endothelialized vessels facilitate patent blood flow *in vivo* and therefore are critical to survival of implanted bioengineered organs by contributing to the delivery of oxygen and nutrients to parenchymal cells (Pellegata AF et al., 2018). We developed our approach to re-endothelialize clinically relevant decellularized porcine scaffolds based on a panel of criteria: 1) adequate histological vascular coverage with endothelial cells, 2) demonstrated increases in cellular metabolism kinetics by non-destructive monitoring to predict patency, and 3) functional hemocompatibility for sustained *in vivo* perfusion. All three criteria were used to refine the methods for endothelial cell seeding, recellularized scaffold perfusion culture, and systematic hemodynamic perfusion to ultimately reach predictive outcomes. Similar to our previous finding that a 10 mg/hour minimum GCR for adequate histological endothelial cell coverage and contrast perfusion in recellularized porcine livers (Mao SA et al., 2017), endothelialized kidney grafts exceeding a minimum GCR threshold of 20 mg/hour in the present study maintained persistent perfusion without thrombosis upon exposure to blood flow. The combination of histologic vascular coverage and lack of coagulation supports near-complete re-endothelialization of the vasculature. This rigorous approach allowed for predictability of the readiness of recellularized kidney grafts for transplantation based on non-destructive evaluation (Uzarski JS et al., 2017).

Differences in human and porcine anatomy required the surgical approach to kidney transplantation used in this study to differ from the standard retroperitoneal access used clinically (Golriz M et al., 2012). While human patients are typically positioned supine, retroperitoneal access in the pig requires the animal to be positioned on its side, reducing operating space in the surgical field and making vascular anastomosis challenging. In this study, midline laparotomy performed on supine pigs provided a larger surgical field but allowed positional changes in transplanted grafts after animal recovery that caused the anastomosed vessels to kink or twist, leading to turbulent hemodynamics near the anastomoses. These complications resulted in thrombosis in the renal artery and/or vein, a major source of graft failure in this study. Thrombotic occlusion was mitigated by securing the kidney graft to the abdominal wall using the native peritoneum that stabilized the graft after recovery, thereby prolonging graft patency. In addition, clopidogrel was added to the protocol (starting on post-operative day 1) to specifically reduce the thrombosis observed at the anastomosis sites after an initial 2 kidneys demonstrated continuous perfusion through 1 or 3 days in the absence of anti-coagulation therapy before ultimately succumbing to thrombosis at the anastomoses. Further improvement of the surgical model to overcome these challenges will facilitate heterotopic kidney transplantation in the retroperitoneum, the standard clinical practice.

Transplanted porcine kidney grafts re-endothelialized with HUVECs maintained vascular patency for up to 1 week post-operatively following *in vivo* orthotopic implantation. Angiographies from the endothelialized grafts performed at regular follow-up intervals showed that patent grafts maintained consistent and uniform perfusion based on gross qualitative visualization (Figure 6; Supplementary Movies 2-4). Despite administration of steroid immunosuppression therapy, a gradual reduction in HUVEC coverage was observed over time, likely due to an immune response to the human cells despite steroid treatment. Considering the continued graft patency based on angiography which was assessed both qualitatively in real-time and quantitatively post-mortem, it was unexpected that most blood vessels would lose human endothelial cell coverage after 7 days. Initial vascular thromboresistance was conferred by endothelial cells seeded in the grafts as decellularized kidney grafts showed thrombosis within minutes despite the direct use of heparin. While the addition of clopidogrel may have provided some thromboresistance, three initial kidney grafts implanted in animals that did not receive clopidogrel were patent until days 1, 3, and 7 before kinking and/or thrombosis at the anastomosis site, indicating that endothelialized kidneys were capable of sustaining renal blood flow.

The finding that a bioengineered kidney can remain patent during endothelial coverage turnover is potentially promising as the immune system is likely to target endothelial cells even with robust immune suppression strategies, and the vasculature can tolerate exposure of the extracellular matrix until native host endothelial cells are deposited into the graft. The continued patency despite loss of HUVECs from the vasculature suggests a possible novel means for maintenance of patency for the bioengineered kidney grafts. We hypothesize that either the HUVECs are being masked in the implants and are not being detected by immunostaining or that the recellularization with HUVECs leads to an anti-thrombogenic surface following the loss of HUVECs, which lends the surface to gradual revascularization by host endothelial cells. Further studies are needed to understand these results. Further work is also needed to stabilize glomerular capillary endothelialization *in vivo* through the recellularization of the epithelial niche and characterization of fenestrations critical to renal function (Ballermann BJ, 2005). These fenestrations, which HUVECs have been demonstrated to be capable of forming *in vitro,* are critical for glomerular filtration (Hamilton RD et al., 2007).

Future research is also needed to develop a better immunosuppression protocol for this porcine model to mitigate xenogenic cell rejection. It should be noted that the loss of HUVECs is a limitation of placing human cells into an animal model (a reverse xenotransplant) and that this would not be indicative of what would occur when decellularized porcine kidney grafts recellularized with HUVECs is transplanted into a human.

Although the present study demonstrated the maintenance of graft patency, further studies are needed to assess and document glomerular function and urine production. Looking ahead to future potential clinical translation of this approach, a short-term strategy would be to source human cells from donor kidneys that were unable to be placed for transplant (Abdelwahab Elhamahmi D et al., 2019). Immunosuppression will still be required for tolerance of bioengineered kidneys produced using this strategy, but the availability of kidneys for transplantation can be increased above current levels. The long-term strategy is to circumvent the need for patient immunosuppression by obtaining autologous cells from the intended transplant recipient using biopsy and/or cellular reprogramming via induced pluripotency/directed differentiation. Recently established protocols detailing directed human induced pluripotent stem cell differentiation into podocytes (Yoshimura Y, et al., 2019), nephron progenitor (Morizane R et al., 2017; Takasato M et al., 2016), and ureteric bud collecting duct cells support this approach (Mae SR, et al., 2019). However, considerable development and scale-up work is needed to produce billions of specialized kidney cells required for scaffold recellularization.

## 5 Conclusion

In summary, this study is the first to demonstrate that human-scale bioengineered kidney grafts developed via perfusion decellularization and subsequent re-endothelialization with HUVECs can remain patent with consistent blood flow for up to 7 days *in vivo*. These results provide a basis for further exploratory studies to assess the potential for using bioengineered kidneys as an alternative to human allograft kidneys.

## Data Availability Statement

The data that support the findings of this study are available from the corresponding author, JR, upon reasonable request.

## Ethics Statement

The present study was carried out in the facilities of American Preclinical Services (Minneapolis, MN), an Association for Assessment and Accreditation of Laboratory Animal Care (AAALAC) approved facility. The study protocol was reviewed and approved by the facility’s Institutional Animal Care and Use Committee (IACUC).

## Author Contributions

JSU, ECB, and EER: performed all in vitro experiments. JSU, ECB, EER, EJV, DSD, and JJR: wrote the paper. MLH, VW, DA, RS, and SSF: assisted with oversight and the development of the surgical model. DSD, SSF and JJR: contributed to experimental planning and oversight. All authors read and approved the manuscript.

## Acknowledgments

We would like to thank Christina Gross, Liisa Carter, and the surgical staff at American Preclinical Services, Inc. (Coon Rapids, MN) for their intellectual contributions to the orthotopic kidney transplantation model. We thank Dr. Brett Anderson for contributing the perfusion bioreactor diagram. We acknowledge Scientific Solutions, LLC for histological tissue processing.

## References

Abdelwahab Elhamahmi, D., Chaly, T., Jr, Wei, G., and Hall, I. E. (2018). Kidney Discard Rates in the United States During American Transplant Congress Meetings. Transplantation Direct, 5(1), e412. https://doi.org/10.1097/TXD.0000000000000849

Abolbashari, M., Agcaoili, S. M., Lee, M. K., Ko, I. K., Aboushwareb, T., Jackson, J. D., Yoo, J. J., and Atala, A. (2016). Repopulation of porcine kidney scaffold using porcine primary renal cells. Acta Biomaterialia, 29, 52–61. https://doi.org/10.1016/j.actbio.2015.11.026

Balleisen, L., and Rauterberg, J. (1980). Platelet activation by basement membrane collagens. Thrombosis Research, 18(5), 725–732. https://doi.org/10.1016/0049-3848(80)90227-3

Ballermann B. J. (2005). Glomerular endothelial cell differentiation. Kidney international, 67(5), 1668–1671. https://doi.org/10.1111/j.1523-1755.2005.00260.x

Bonandrini, B., Figliuzzi, M., Papadimou, E., Morigi, M., Perico, N., Casiraghi, F., et al. (2014). Recellularization of well-preserved acellular kidney scaffold using embryonic stem cells. Tissue Engineering. Part A, 20(9-10), 1486–1498. https://doi.org/10.1089/ten.TEA.2013.0269

Caralt, M., Uzarski, J. S., Iacob, S., Obergfell, K. P., Berg, N., Bijonowski, B. M., et al. (2015). Optimization and critical evaluation of decellularization strategies to develop renal extracellular matrix scaffolds as biological templates for organ engineering and transplantation. American Journal Of Transplantation: Official Journal of the American Society of Transplantation and the American Society of Transplant Surgeons, 15(1), 64–75. https://doi.org/10.1111/ajt.12999

Gilpin, S. E., and Ott, H. C. (2015). Using nature’s platform to engineer bio-artificial lungs. Annals of the American Thoracic Society, 12 *Suppl 1*, S45–S49. https://doi.org/10.1513/AnnalsATS.201408-366MG

Goh, S. K., Bertera, S., Olsen, P., Candiello, J. E., Halfter, W., Uechi, G., et al. (2013). Perfusion-decellularized pancreas as a natural 3D scaffold for pancreatic tissue and whole organ engineering. Biomaterials, 34(28), 6760–6772. https://doi.org/10.1016/j.biomaterials.2013.05.066

Golriz, M., Fonouni, H., Nickkholgh, A., Hafezi, M., Garoussi, C., and Mehrabi, A. (2012). Pig kidney transplantation: an up-to-date guideline. European Surgical Research. Europaische Chirurgische Forschung. Recherches Chirurgicales Europeennes, 49(3-4), 121–129. https://doi.org/10.1159/000343132

Hamilton, R. D., Foss, A. J., and Leach, L. (2007). Establishment of a human in vitro model of the outer blood-retinal barrier. Journal of Anatomy, 211(6), 707–716. https://doi.org/10.1111/j.1469-7580.2007.00812.x

Lentine, K. L., Smith, J. M., Hart, A., Miller, J., Skeans, M. A., Larkin, L., et al. (2022). OPTN/SRTR 2020 Annual Data Report: Kidney. American Journal of transplantation: Official Journal of the American Society of Transplantation and the American Society of Transplant Surgeons, 22 *Suppl 2*, 21–136. https://doi.org/10.1111/ajt.16982 Supplementary Material

Leuning, D. G., Witjas, F. M. R., Maanaoui, M., de Graaf, A. M. A., Lievers, E., Geuens, T., et al. (2019). Vascular bioengineering of scaffolds derived from human discarded transplant kidneys using human pluripotent stem cell-derived endothelium. American Journal of Transplantation: OfficialJjournal of the American Society of Transplantation and the American Society of Transplant Surgeons, 19(5), 1328–1343. https://doi.org/10.1111/ajt.15200

Mae, S. I., Ryosaka, M., and Osafune, K. (2019). Protocol to Generate Ureteric Bud Structures from Human iPS Cells. Methods in Molecular Biology (Clifton, N.J.), 1926, 117–123. https://doi.org/10.1007/978-1-4939-9021-4_10

Mao, S., Glorioso, J., Elgilani, F., De Lorenzo, S., and Deeds. M. (2017). Sustained in vivo perfusion of a re-endothelialized tissue engineered porcine liver. Int J Transplant Res Med, 3, 031.

Miyoshi, T., Hiratsuka, K., Saiz, E. G., and Morizane, R. (2020). Kidney organoids in translational medicine: Disease modeling and regenerative medicine. Developmental dynamics: An Official Publication of the American Association of Anatomists, 249(1), 34–45. https://doi.org/10.1002/dvdy.22

Molema, G., and Aird, W. C. (2012). Vascular heterogeneity in the kidney. Seminars in nephrology, 32(2), 145–155. https://doi.org/10.1016/j.semnephrol.2012.02.001

Morizane, R., and Bonventre, J. V. (2017). Generation of nephron progenitor cells and kidney organoids from human pluripotent stem cells. Nature Protocols, 12(1), 195–207. https://doi.org/10.1038/nprot.2016.170

Nakayama, K. H., Batchelder, C. A., Lee, C. I., and Tarantal, A. F. (2010). Decellularized rhesus monkey kidney as a three-dimensional scaffold for renal tissue engineering. Tissue Engineering. Part A, 16(7), 2207–2216. https://doi.org/10.1089/ten.tea.2009.0602

Orlando, G., Farney, A. C., Iskandar, S. S., Mirmalek-Sani, S. H., Sullivan, D. C., Moran, E., et al. (2012). Production and implantation of renal extracellular matrix scaffolds from porcine kidneys as a platform for renal bioengineering investigations. Annals of Surgery, 256(2), 363–370. https://doi.org/10.1097/SLA.0b013e31825a02ab

Ott, H. C., Clippinger, B., Conrad, C., Schuetz, C., Pomerantseva, I., Ikonomou, L., et al. (2010). Regeneration and orthotopic transplantation of a bioartificial lung. Nature Medicine, 16(8), 927–933. https://doi.org/10.1038/nm.2193

Ott, H. C., Matthiesen, T. S., Goh, S. K., Black, L. D., Kren, S. M., Netoff, T. I., et al. (2008). Perfusion-decellularized matrix: using nature’s platform to engineer a bioartificial heart. Nature Medicine, 14(2), 213–221. https://doi.org/10.1038/nm1684

Pellegata, A. F., Tedeschi, A. M., and De Coppi, P. (2018). Whole Organ Tissue Vascularization: Engineering the Tree to Develop the Fruits. Frontiers in Bioengineering and Biotechnology, 6, 56. https://doi.org/10.3389/fbioe.2018.00056

Peloso, A., Ferrario, J., Maiga, B., Benzoni, I., Bianco, C., Citro, A., et al. (2015). Creation and implantation of acellular rat renal ECM-based scaffolds. Organogenesis, 11(2), 58–74. https://doi.org/10.1080/15476278.2015.1072661

Petersen, T. H., Calle, E. A., Zhao, L., Lee, E. J., Gui, L., Raredon, M. B., et al. (2010). Tissue-engineered lungs for in vivo implantation. Science (New York, N.Y.), 329(5991), 538–541. https://doi.org/10.1126/science.1189345

Ross, E. A., Williams, M. J., Hamazaki, T., Terada, N., Clapp, W. L., Adin, C., et al (2009). Embryonic stem cells proliferate and differentiate when seeded into kidney scaffolds. Journal of the American Society of Nephrology: JASN, 20(11), 2338–2347. https://doi.org/10.1681/ASN.2008111196

Shaheen, M. F., Joo, D. J., Ross, J. J., Anderson, B. D., Chen, H. S., Huebert, R. C., Li, Y., Amiot, B., Young, A., Zlochiver, V., Nelson, E., Mounajjed, T., Dietz, A. B., Michalak, G., Steiner, B. G., Davidow, D. S., Paradise, C. R., van Wijnen, A. J., Shah, V. H., Liu, M., … Nyberg, S. L. (2020). Sustained perfusion of revascularized bioengineered livers heterotopically transplanted into immunosuppressed pigs. Nature Biomedical Engineering, 4(4), 437–445. https://doi.org/10.1038/s41551-019-0460-x

Song, J. J., Guyette, J. P., Gilpin, S. E., Gonzalez, G., Vacanti, J. P., and Ott, H. C. (2013). Regeneration and experimental orthotopic transplantation of a bioengineered kidney. Nature Medicine, 19(5), 646–651. https://doi.org/10.1038/nm.3154

Takasato, M., Er, P. X., Chiu, H. S., and Little, M. H. (2016). Generation of kidney organoids from human pluripotent stem cells. Nature Protocols, 11(9), 1681–1692. https://doi.org/10.1038/nprot.2016.098

United States Renal Data System. 2022 USRDS Annual Data Report: Epidemiology of Kidney Disease in the United States. National Institutes of Health, National Institute of Diabetes and Digestive and Kidney Diseases, Bethesda, MD, 2022.

Uygun, B. E., Soto-Gutierrez, A., Yagi, H., Izamis, M. L., Guzzardi, M. A., Shulman, C., et al. (2010). Organ reengineering through development of a transplantable recellularized liver graft using decellularized liver matrix. Nature Medicine, 16(7), 814–820. https://doi.org/10.1038/nm.2170

Uzarski, J. S., DiVito, M. D., Wertheim, J. A., and Miller, W. M. (2017). Essential design considerations for the resazurin reduction assay to noninvasively quantify cell expansion within perfused extracellular matrix scaffolds. Biomaterials, 129, 163–175. https://doi.org/10.1016/j.biomaterials.2017.02.015

Uzarski, J. S., Xia, Y., Belmonte, J. C., and Wertheim, J. A. (2014). New strategies in kidney regeneration and tissue engineering. Current opinion in nephrology and hypertension, 23(4), 399–405. https://doi.org/10.1097/01.mnh.0000447019.66970.ea

Wang, Y., Bao, J., Wu, Q., Zhou, Y., Li, Y., Wu, X., et al. (2015). Method for perfusion decellularization of porcine whole liver and kidney for use as a scaffold for clinical-scale bioengineering engrafts. Xenotransplantation, 22(1), 48–61. https://doi.org/10.1111/xen.12141

Willette, R. N., Storer, B. L., Clark, R. K., and Ohlstein, E. H. (1994). Human laminin produces human platelet aggregation in vitro. Life Sciences, 55(5), 379–388. https://doi.org/10.1016/0024-3205(94)00649-0

Wu, H., Uchimura, K., Donnelly, E. L., Kirita, Y., Morris, S. A., and Humphreys, B. D. (2018). Comparative Analysis and Refinement of Human PSC-Derived Kidney Organoid Differentiation with Single-Cell Transcriptomics. Cell Stem Cell, 23(6), 869–881.e8. https://doi.org/10.1016/j.stem.2018.10.010

Yoshimura, Y., Taguchi, A., Tanigawa, S., Yatsuda, J., Kamba, T., Takahashi, S., et al. (2019). Manipulation of nephron-patterning signals enables selective induction of podocytes from human pluripotent stem cells. Journal of the American Society of Nephrology, 30(2), 304–321. https://doi.org/10.1681/ASN.2018070747

Yu, Y. L., Shao, Y. K., Ding, Y. Q., Lin, K. Z., Chen, B., Zhang, H. Z., et al. (2014). Decellularized kidney scaffold-mediated renal regeneration. Biomaterials, 35(25), 6822–6828. https://doi.org/10.1016/j.biomaterials.2014.04.074

Zambon, J. P., Ko, I. K., Abolbashari, M., Huling, J., Clouse, C., Kim, T. H., et al. (2018). Comparative analysis of two porcine kidney decellularization methods for maintenance of functional vascular architectures. Acta Biomaterialia, 75, 226–234. https://doi.org/10.1016/j.actbio.2018.06.004

