## Supplementary Materials for "Sustained In Vivo Perfusion of a Re-Endothelialized Tissue Engineered Kidney Graft in a Human-Scale Animal Model"

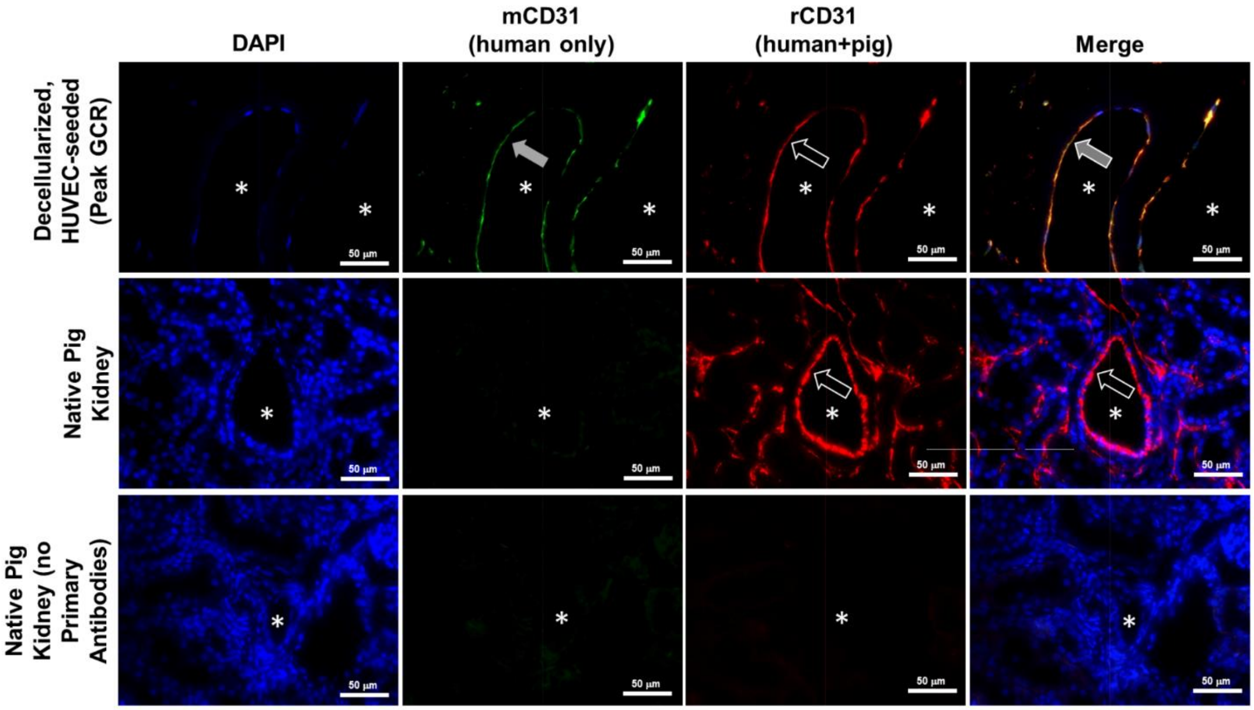


**Supplementary Figure S1. Characterization of differential CD31 antibody reactivity for identification of human or pig endothelial cells in chronically transplanted kidney grafts.** Endothelial cells were identified as donor (human) or recipient (porcine) in explanted kidney grafts using immunofluorescence staining with two different CD31 antibodies. A decellularized kidney containing only human cells and a native kidney containing only porcine cells were used to demonstrated species reactivity in two antibodies. Grey arrow fill indicates positive expression by mouse CD31 antibody (mCD31; green stain), which reacts positively only in human endothelial cells. White arrow outline indicates positive reactivity by rabbit CD31 antibody (rCD31; red stain), which reacts to both human and pig endothelial cells. Orange color shows colocalization of both antibodies, indicating human endothelial cells. Asterisks denote the lumens of patent blood vessels. Scale bars, 50 µm.

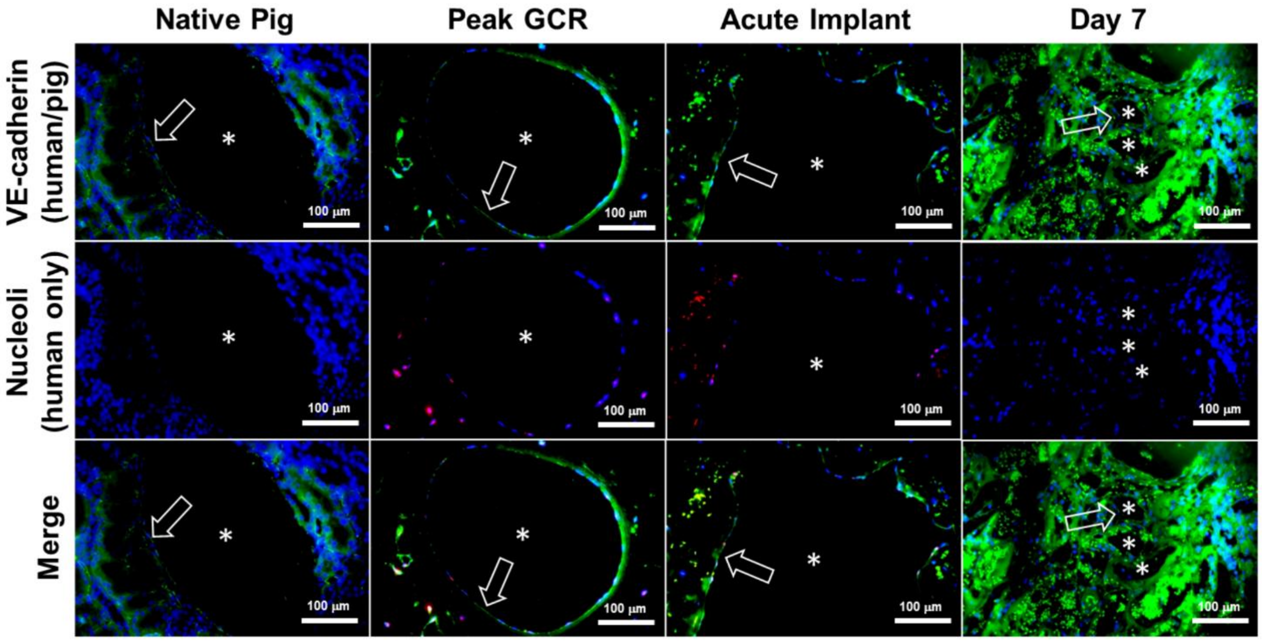


**Supplementary Figure S2: Characterization of donor human endothelial cell loss and replacement with recipient porcine endothelial cells in chronically transplanted endothelialized kidney grafts.** Patent endothelialized kidney grafts were explanted at regular follow-up time intervals following chronic orthotopic transplantation in a pig model. Native pig kidneys, explanted grafts, or non-implanted and HUVEC-recellularized kidneys at Peak GCR were flushed with saline, fixed, and processed for immunofluorescence staining to identify human endothelial cells in the vasculature using an anti-human nucleoli antibody (red stain). Asterisks denote the lumens of patent blood vessels. White arrow outline indicates positive reactivity by VE-cadherin antibody (green stain), which reacts to both human and pig endothelial cells. Top row: VE-cadherin/DAPI overlay. Middle row: Nucleoli/DAPI overlay. Bottom row: VE-cadherin/Nucleoli/DAPI merged images. Note that the presence of residual blood causes background staining in some images. Donor human endothelial cells present in Peak GCR controls and Acute Implant grafts were completely absent from the graft by day 7, while new pig endothelialization started to occur at day 7. Scale bars, 100 µm.
